# No evidence that visual impulses enhance the readout of retrieved long-term memory contents from EEG activity

**DOI:** 10.1101/2024.04.19.590215

**Authors:** Sander van Bree, Abbie Sarah Mackenzie, Maria Wimber

## Abstract

The application of multivariate pattern analysis (MVPA) to electroencephalography (EEG) data allows neuroscientists to track neural representations at temporally fine-grained scales. This approach has been leveraged to study the locus and evolution of long-term memory contents in the brain, but a limiting factor is that decoding performance remains low. A key reason for this is that processes like encoding and retrieval are intrinsically dynamic across trials and participants, and this runs in tension with MVPA and other techniques that rely on consistently unfolding neural codes to generate predictions about memory contents. The presentation of visually perturbing stimuli may experimentally regularize brain dynamics, making neural codes more stable across measurements to enhance representational readouts. Such enhancements, which have repeatedly been demonstrated in working memory contexts, remain to our knowledge unexplored in long-term memory tasks. In this study, we evaluated whether visual perturbations—or *pings*—improve our ability to predict the category of retrieved images from EEG activity during cued recall. Overall, our findings suggest that while pings evoked a prominent neural response, they did not reliably produce improvements in MVPA-based classification across several analyses. We discuss possibilities that could explain these results, including the role of experimental and analysis parameter choices and mechanistic differences between working and long-term memory.

## Introduction

A central question in memory research is how the brain retrieves information stored in long-term memory (LTM) in the service of adaptive behaviour. This research topic has inspired work from a variety of angles, involving different experimental protocols and methods—including neuroimaging modalities. Electroencephalography (EEG) and magnetoencephalography (MEG) have proven an integral part of this project because they capture brain dynamics on a sub-second resolution. Such granularity is crucial, given that memory retrieval typically unfolds on the order of seconds, with the neural cascades underpinning memory retrieval evolving even faster (Staresina & Wimber, 2019).

To study the evolution of retrieved contents in the brain, one widely pursued family of techniques is multivariate pattern analysis (MVPA)—more broadly known as classification or decoding (Haxby et al., 2014; Grootswagers et al., 2017). These tools extract and upweight signal dimensions that robustly covary with retrieved memory contents, effectively boosting the signal-to-noise ratio of associated neural activity. MVPA has been successfully used to enrich our understanding of memory, including how information is encoded (Fritch et al., 2020; Kragel et al., 2017; Kuhl et al., 2012), consolidated (Deuker et al., 2013; Maguire, 2014; Schreiner et al., 2021), and reinstated during memory recall (i.e., pattern completion; Danker & Anderson, 2010; Favila et al., 2020; Rissman & Wagner, 2012; Xue, 2018).

Despite such advancements, the decoding of long-term memory contents in electrophysiology data typically remains only slightly above chance, impairing our ability to study the evolution of neural patterns of interest. One reason for this limitation is that memory processes and their associated brain activity are highly dynamic, which results in variable patterns across trials and participants (ter Wal et al., 2021; Madore & Wagner, 2022). Indeed, MVPA and most other EEG-based analyses rely for their robust predictions on the existence of a detectably constant cascade of neural patterns across measurements (van Bree et al., 2022). This clash between variability in neural processes on the one hand and the constancy assumption of our analyses on the other may cause us to miss representations of interest, or to obtain different results depending on what experimental event we timelock EEG data to (e.g., retrieval cues vs button presses; Linde-Domingo et al., 2019). A factor that further hampers our ability to robustly decode representations is that retrieval comes with fainter neural patterns to begin with compared to perception (Favila et al., 2020; Pearson et al., 2015; Favila et al., 2022). Together, these points invite creative techniques that improve our ability to infer long-term memory representations from dynamic brain activity.

In this study, we explore a perturbational method that has the potential to mitigate two issues at the same time: low signal fidelity at the level of measurement, and variability in neural processing dynamics. Specifically, in this EEG study we evaluated whether the presentation of a high contrast visual stimulus—henceforth referred to as a “ping”—during LTM retrieval enhances the readout of signatures of retrieved content. In motivating the hypothesis that *pings* boost the decodability of LTM representations, we built directly onto recent successful efforts in the domain of working memory (WM). In that context, pings have been used to enhance the decodability of the orientation (Wolff et al., 2015, 2017, 2020; Ten Oever et al., 2020; Yang et al., 2023) and colour (Kandemir et al., 2023) of objects actively maintained in WM, as well as anticipated target locations (Duncan et al., 2023). A preliminary explanation for these findings is that pings induce a robust evoked response that interacts and indeed boosts the footprint of active neural representations, enhancing their SNR (Barbosa et al., 2021). Specifically, pings may regularize neural dynamics across trials and participants by producing a phase reset of brain oscillations that coordinate information processing across neuronal populations. In support of this, visual stimuli presented during memory tasks have been shown to reset the phase of low-frequency brain oscillations that are implicated in encoding and retrieval (Rizzuto et al., 2003; Haque et al., 2015; audiovisual stimuli in Cruzat et al., 2021). Thus, by inducing pings at experimentally controlled moments, researchers may gain a level of control over variability in synchronized activity across information-coding neurons, making their dynamics more similar across measurements to improve the predictive power of MVPA.

Importantly however, while ping-based methods have been shown to work in WM contexts, to our knowledge it has not been explored whether they generalize to LTM research in which information is retrieved from stored representations. The purpose of this study then, is to systematically explore the possibility that pings can enhance the readout of reactivated long-term memory contents. To this end, we presented participants with pings as memory processes were actively engaged during cued recall, evaluating whether retrieved representations are more robustly discernible after ping onset. On the whole, we find no compelling evidence that pings boost the classification of retrieved image pairs from EEG activity.

## Methods

### Participants

We recruited thirty-three volunteers (22 women, M_age_ = 23.8 years, SD_age_ = 2.6 years, range = 18 to 31) with normal or corrected-to-normal vision, and with no history of epileptic attacks or neuropsychological conditions that could interfere with the examined study effects. The sample size required to derive a reliable effect was estimated based on (Wolff et al., 2017), though our estimation was limited by the fact that all previous work was in a WM context. One participant did not finish the experiment because they were unwell, and following data inspection, two participants were removed because of poor data quality due to a large number of high impedance channels, and one because of stimulus trigger issues. Thus, EEG-based analyses were conducted based on 29 participants. For behavioural analyses, the first four participants were excluded because of missing button press triggers, which, with the further exclusion of the participant who did not complete the experiment, resulted in an analysis of 28 participants (participants with noisy EEG data were included in the behavioural analysis).

Participants were informed about the details of the experiment in advance—including its duration, protocol, and methods—but were left naïve with respect to the purpose and hypotheses associated with the presentation of visual pings. Participants provided their written consent, and after the experiment, they were debriefed and given information about the central manipulation and hypothesis upon request, and they were compensated for their time with £9 per volunteered hour. The study was approved by the Ethical committee of the College of Science and Engineering of the University of Glasgow (Application number: 300210113).

### Stimulus and apparatus

The presentation of stimuli was controlled using PsychoPy (version 2021.2.3; Peirce et al., 2019) running on Windows 10. Stimuli were presented on a CRT monitor (53.3 cm; 1024 by 768 pixels) operating at a refresh rate of 60 Hz. Participants were seated in a magnetically shielded room in a chinrest 65 cm from the screen, or at an approximately similar distance from the screen outside the chinrest if they experienced discomfort. Throughout the experiment, a fixation cross (with a visual angle of 0.44°) was presented in the centre of a constantly presented grey background (RGB = 128 128 128; PsychoPy default). All centrally presented stimuli overrode the fixation dot. The visual impulse (i.e., ping) was a single full-contrast bullseye stimulus presented at the centre of the screen for 200 milliseconds (ms; with a diameter of 13° and 0.31° cycles per degree). The ping was generated using MATLAB and edited using GIMP (GNU Image Manipulation Program version 2.10.32).

In the main memory task, participants learned associations between action verbs and images, and were later prompted with the action verb to retrieve the associated image. The action verbs were selected based on usage frequency (largely based on Linde-Domingo et al., 2019) and the image stimulus set was a combination of 192 colour images collated across various royalty free databases, including the Bank of Standardized Stimuli (BOSS, Brodeur et al., 2010), and the SUN database (Xiao et al., 2010). The selected 192 images were constructed to follow a nested category structure of three embedded hierarchical levels. At the top level, the set consisted of 96 objects and 96 scenes, which were in turn composed at the middle level of 48 animate and 48 inanimate objects and 48 indoor and 48 outdoor scenes. Moving down to the bottom level, each of the middle level categories branched out into 4 categories (e.g., for animate objects: birds, insects, mammals, and marine animals), each of which contained 12 specific instances (e.g., twelve specific birds). We chose this nested hierarchy of stimulus categories because we did not know a priori what dimension of retrieved memories would be effectively decodable, so we included multiple levels of abstraction and chose one level based on pre-defined criteria (See *Level Selection*). The objects were presented on a white square matching in size to scene images (i.e., the visual degrees of all stimulus categories were 13°). Key presses were registered using a standard QWERTY keyboard.

### Procedure

The main experiment consisted of 8 blocks, each with an encoding, distractor, recall, and recognition phase (Fig. 1A). In total, the main experiment lasted between approximately 45 and 65 minutes depending on the duration of self-paced breaks and electrode impedance maintenance. Before the main experiment, participants were provided with a practice run that covered each phase using example verbs and images that were not used in the main experiment. A standardized set of verbal instructions were provided to guide participants through the practice run. If the participant reported not understanding the task or if they did not give accurate responses, the practice run and instructions were repeated. Then, the main experiment commenced, throughout which EEG was acquired. At the start of each experimental phase, a screen was presented with a reminder of the task instructions and required response keys.

**Figure 1.**
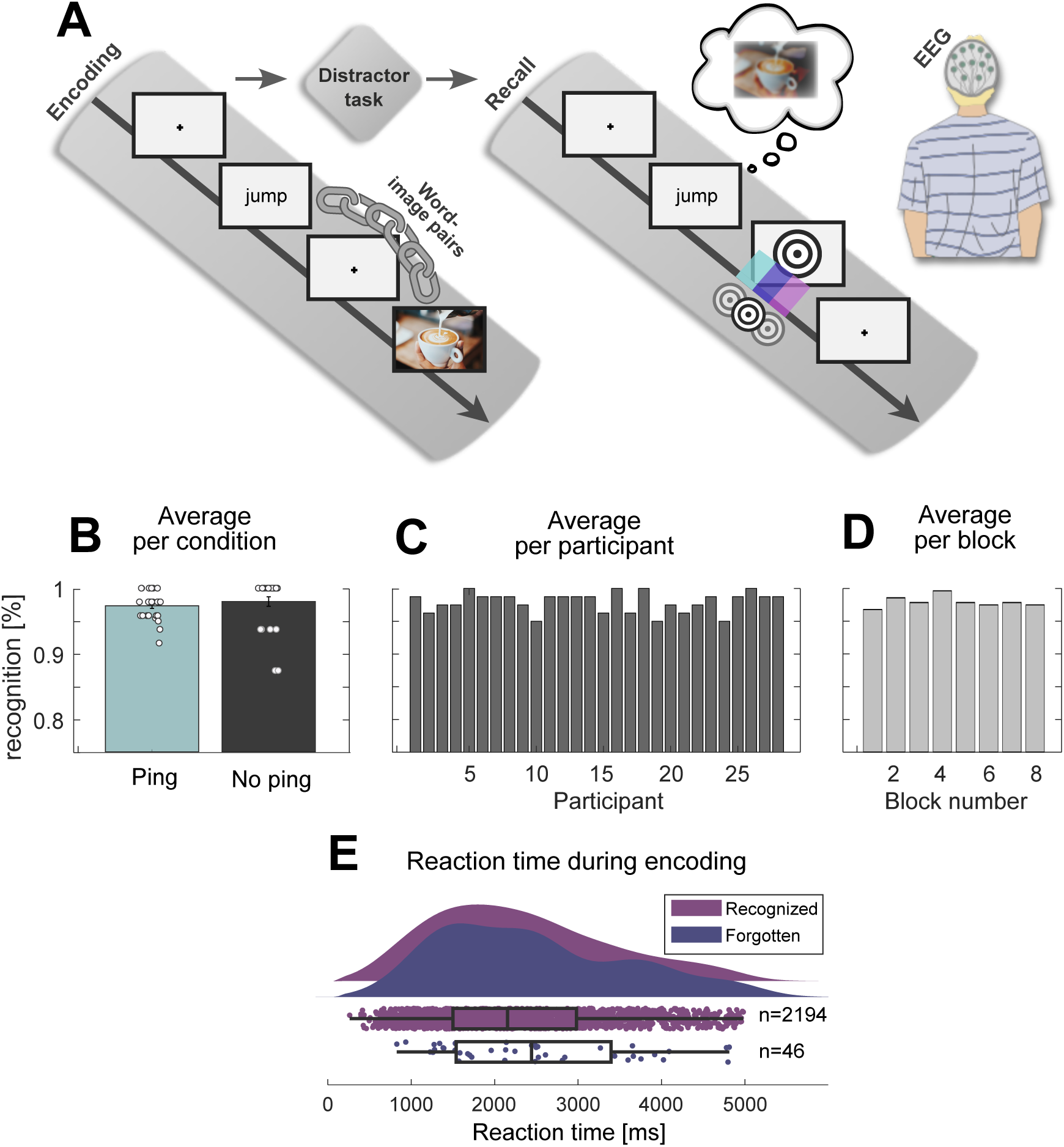
Paradigm and behavioural results. **(A)** Experimental paradigm. The encoding phase consisted of a word-image pair learning task. This was followed by a distractor task intended to wash out working memory effects. Then, during the critical recall phase, participants were cued with words to retrieve the paired image while visual perturbations (pings) were presented in 75% of trials. In a fourth phase, recognition performance was tested (not displayed). **(B)** Average performance during the recognition task for trials with and without pings, collapsing across blocks for each participant. Datapoints are individual participants. **(C)** Average recognition performance per participant (i.e., collapsing blocks). **(D)** Average recognition performance per block (i.e., collapsing participants). **(E)** Average reaction time during encoding for subsequently recognized and forgotten trials, collapsing across blocks. Note: in B, C, and D, the y-axis is truncated due to high recognition performance.

In the encoding phase, participants learned to build a mental association between action verbs and paired images. First, a verb was presented for 1500 ms (white, *OpenSans* font). Then, after 1000 ms, the associated image was presented until the spacebar was pressed to indicate the association was encoded (with a 6000 ms limit). Then, after a 1000 ms delay, the next verb was presented. During each block’s encoding phase, 10 unique verb-image pairs were learned in one shot. This resulted in 80 encoded pairs across the full experiment, with the images pseudo-randomly selected from the full stimulus set such as to maintain an equal distribution of top-level stimulus categories (40 objects and 40 scenes) and fully random selection over nested middle and bottom levels for each ping and no-ping condition.

The distractor phase that followed was included to flush out WM effects. Here, participants performed an odd-even task lasting 20 seconds. A number between 1 and 99 was presented in the centre of the screen (white, *OpenSans* font), and participants were instructed to press left key for odd numbers, and right key for even numbers. Following a left or right key press, the next number was presented immediately. Participants’ average performance was displayed at the end of the distractor phase, marked as the proportion of correct responses. This data was not further analysed.

Next in each block, the recall phase tested our central manipulation of a ping-based visual perturbation. In this phase, participants recalled the learned verb-image associations of the encoding phase. First, one of the ten encoded verbs was presented for 2000 ms, serving as the retrieval cue that prompted recall of the associated image. In 75% of trials, a visual impulse was presented in either of three time bins: between 500 to 833.33 ms (“early ping”), 833.34 to 1116.67 ms (“middle ping”), or 1116.68 to 1500 ms (“late ping”) after the onset of the retrieval cue, with a uniform distribution of possible ping times within each bin. This window was chosen on the basis that previous research on cued recall paradigms suggests this is the moment of maximum memory reinstatement (Staresina & Wimber, 2019). In 25% of trials, no visual impulse was presented in order to derive a baseline for statistical hypothesis testing. Participants pressed the left key to indicate that they had forgotten the image associated with the verb cue, or right key to indicate they remembered it. Key presses only resulted in a new trial after 1700 ms following retrieval cue onset (i.e., 200 ms after the latest possible ping). With presses earlier than that, nothing happened. Participants were given a visual indication that key presses were available via disappearance of the retrieval cue (at its offset; 2000 ms). During the recall phase, each of the 10 encoded verb-image pairs were tested four times, resulting in 40 recall trials per block, and 320 trials in total, comprising 160 objects and 160 scenes. Within participants, each of the four conditions—early, middle, late, and no ping—were configured to present object and scene images equally often (i.e., the top-level stimulus category), with the nested mid and bottom-level categories randomized. The sequence of presented stimulus level categories, pinging conditions, and verb-image pairs was fully randomized within and across blocks to mitigate order effects. For the within block randomization, while the 40 recall trials were fully randomized, we ensured the same pair was never recalled twice in direct succession.

Finally, since the cued recall phase only included subjective memory judgments, a recognition phase was included to obtain an objective measure of memory performance for the verb-image pairs. During this two-alternative forced choice task, one of the 10 encoded verbs was presented in the centre of the screen, with two images (visual angle of 7.8°) presented underneath, one on the left-hand, and one on the right-hand side of the screen. Participants chose which of the two images was paired with the central action verb using a left or right key press (with a 5000 ms time limit). The location of the correctly paired image was randomized between the left and right location. The lure image was always another old image from the immediately preceding encoding phase. Each of the 10 encoded verb-image pairs was tested once in a random sequence. Note that we designed this study to expend most of the available study time on the recall phase to maximize the statistical power of our main analysis, with the recognition phase serving chiefly as a basic check to ensure participants were not skipping through the experiment without memorizing verb-image pairs.

### EEG acquisition and preprocessing

The data was recorded using a 64-channel passive EEG BrainVision system (BrainAmp MR; Brain Products) with a sampling rate of 1000 Hz. For our recording software we used BrainVision Recorder (Brain Products). The 64 Ag/AgCl electrodes were positioned in accordance with the extended international 10-20 system. Due to a necessary change in the recording system, a different EEG cap type (EasyCap) was used for participants 1 to 14 (subset 1) and 15 to 33 (subset 2). In the first subset, the ground electrode was located on the back of the head, below occipital electrode Oz, and two EOG channels were used to monitor eye movements (placed below and next to the eye; VEOG and HEOG). In the second subset, the ground electrode was on the midline frontal location AFz, and one EOG channel was used to measure eye movements (placed below the eye; VEOG). Furthermore, the cap used in the second subset included channels FT9 and FT10. For event related potential analyses, we included only electrodes common to both caps to enable a universal visualization of brain activity. Most electrode impedances were kept below 25 kiloΩ, and electrodes with outlier impedances were removed during preprocessing, with their associated data interpolated (see below).

Preprocessing was performed using FieldTrip (Oostenveld et al., 2011) in MATLAB (the MathWorks). First, the continuous EEG data was split up into two datasets: one with all trials epoched relative to retrieval cues, and one with trials epoched relative to pings and no-ping (defined by randomly sampling ping times of the pinged trials, yielding so-called “pseudo-pings”). Put differently, the data was locked once to *t* = 0 defined as the retrieval cue, and once to *t* = 0 defined as the manipulation of interest or a baseline alternative. In both cases, the epoched trials were 4 seconds in duration (-1 to 3 seconds relative to the event of interest).

Each dataset was filtered between 0.05 and 80 Hz and downsampled to 250 Hz. Next, bad trials and channels with outlier impedance levels were manually removed via visual inspection. Subsequently, eye movement and muscle artefacts were identified and removed using ICA decomposition, and removed channels were interpolated using spline interpolation (with the FieldTrip function *ft_scalpcurrentdensity*). Finally, the data was re-referenced using a common average and a Laplacian method (current source density), deriving separate data structures for cue-locked and ping-locked analyses.

### Behavioural analysis

The experiment was designed to result in high or even ceiling memory performance in order to obtain a maximal number of successfully remembered trials, and to optimally evaluate the central hypothesis of a ping-induced decodability enhancement. We report objective performance for the memory test conducted in the recognition phase, both across pinging conditions (Fig. 1B), participants (Fig. 1C), and across blocks (Fig. 1D). We also report subjective judgments during the recall phase, quantifying how often participants report remembering versus forgetting the word-image pair. Reaction time (RT) during the recall phase is uninformative, because as described in the Procedure section, the response key was locked until 1700 ms after cue onset, at which point participants likely had already retrieved the associated image (Staresina & Wimber, 2019). Indeed, participants reported actively waiting for response buttons to become available. Thus, we instead analysed RT during the encoding phase as a function of whether the word-image pair was subsequently recognized or not. These RT data were collapsed across participants and blocks (Fig. 1E). For the proceeding analyses, both subsequently recognized and forgotten trials were included.

### ERP Analysis

For the ERP analyses, only channels common to both electrode cap subsets were used. We applied two types of ERP analyses, one locked to (pseudo-)pings and one to retrieval cues. FieldTrip was used to downsample the data to 250 Hz and a band-pass filter between 0.2 and 40 Hz was used. The data was baseline-corrected from -200 ms to 0 ms from events of interest. For ERP traces, we calculated the average activity across posterior channels (C3, C4, P3, P4, O1, O2, Cz, Pz, Oz, CP1, CP2, C1, C2, P1, P2, CP3, CP4, PO3, PO4, PO7, PO8, CPz, POz). For ERP topographies, we used the 61 channels common to both ERP cap types. We statistically evaluated whether pings resulted in higher amplitude ERPs compared to no-ping trials using non-parametric Monte Carlo permutation tests applied to each channel, correcting for multiple comparisons using Bonferroni correction as implemented in FieldTrip, averaging activity from 200 to 400 ms after pseudo-pings (alpha = 0.05; 10^5^ randomizations).

### MVPA analysis

For MVPA, all EEG channels available per electrode cap type were used except EOG channels. Depending on the analysis, we trained and tested either a multi-class LDA using FieldTrip (*ft_timelockstatistics*), or a binary-class LDA using the MVPA Light toolbox (Treder, 2020). We classified EEG data re-referenced using a Laplacian transform on the basis that it accentuates local patterns (Kayser & Tenke, 2015). All classifier analyses were performed on the recall phase, where our main hypothesis could be evaluated. Unless specified otherwise, analyses were carried out on the retrieval cue-locked dataset. We downsampled the data from 250 Hz to 50 Hz by applying a moving average with a window length of 140 ms, moving in steps of 20 ms. During each step, a Gaussian-weighted mean was applied in which the centre data sample of the window was multiplied by 1, and the tail samples by 0.15 (FWHM = ∼81 ms). In a subsequent step, sample by sample, the data was z-scored across channels (i.e., setting every channel to mean = 0 and standard deviation = 1), followed by training and testing using LDA. To evaluate decoder performance, we applied k-fold cross validation (5 folds, with 25 repetitions). For binary class decoding, we used area under the receiver operating characteristics curve (AUC) as a performance metric because it adjusts for class imbalances (Grootswagers et al., 2017; Xie & Qiu, 2007). For multi-class decoding, where standard AUC is unavailable, we used accuracy and factored in level-specific differences in chance levels. To infer decoding performance values under the null hypothesis, depending on the analysis, we either used no-ping trials or ping trials with shuffled class labels (100 1^st^-level permutations, each with 3 repetitions). All analyses were restricted to the period before button presses were made (i.e., < 2000 ms).

### Level selection

We used a multi-class LDA on no-ping trials to determine which retrieved stimulus category (top, middle, or bottom level) is most robustly detectable in the data when our main experimental manipulation was not applied. This level was then locked in for subsequent analyses that relate to our key hypothesis of ping-induced decoder enhancement. We selected the level with a high baseline performance to offer a conservative starting point from which we could establish whether pings are a powerful tool to further enhance decodability. However, as we will see in the results, stimulus selection rationales matter minimally because we found no reliable level differences in the no-ping decoder across levels to begin with. For statistics, we performed a Wilcoxon rank sum test comparing the empirical and shuffled decoding performance for each level, in the way described in the next section.

### Main analysis

For the statistical analysis of the main hypothesis, we used two-level permutation testing for the ping versus shuffle decodability comparison, and a Wilcoxon ranked sum test for the ping versus no-ping comparison. The former approach, which is based on van Bree et al., 2022, implemented the following algorithm in pseudo-code—applied window-by-window:

1. For each 2^nd^-level permutation (10^5^ times): Grab one random window-specific decodability value from the 1^st^-level distribution of the 25 permutations of each participant and average the result. This yields 10^5^ permuted averages.
2. Generate one empirical p-value by calculating the percentile of the average empirical decoding value within the distribution of permuted averages.

The latter approach involved taking the Wilcoxon signed-rank test between the distribution of empirical decoder results and 1^st^-level permutation results across participants. We opted for a Wilcoxon test over cluster-based methods because it makes minimal assumptions about the distribution of decoding results (Wilcoxon, 1945; Grootswagers et al., 2017). For both approaches, we adjusted the resulting p-values across windows for their false discovery rate (FDR). Since the p-values are not independent across time, we applied the approach by Benjamini & Yekutieli (2001).

Finally, for ping-locked analyses we restricted statistical analyses between 0 and 500 ms from ping onset. For analyses locked to retrieval cue, we analysed 500 to 2000 ms from cue, which is the approximate range where memory reactivation is maximal (Staresina & Wimber, 2019).

### Condition-relative decoding peaks

In addition to our main analysis, we carried out a presumably more sensitive analysis to evaluate the possibility of ping-induced decoding enhancements. We reasoned that even if visual pings do not offer an enhancement of LTM decoding performance that is strong enough to emerge in a direct ping-to-no ping or ping-to-shuffle comparison, there could still be a weaker effect that is detectable by factoring in the relative order of decoding peaks across pinging conditions. Specifically, we tested whether trials with an early, middle, and late ping tended to have, respectively, earlier, later, and even later decoding performance peaks. In other words, we tested to what extent decoding peaks captured ping presentation orders (see Linde-Domingo et al., 2019; Mirjalili et al., 2021 for similar peak selection approaches).

First, we took every participant’s SOA-specific decoding time series—early, middle, and late—and extracted one peak (specified below). Then, we calculated a *peak order distance* (POD) per participant, defined as the absolute serial distance between the order of extracted peaks and true ping presentation order, given by the formula:

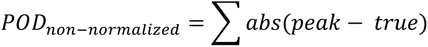

For example, if the decoder peak came first for early ping trials (1 − 1), third for middle pings trials (3 − 2), and second for late ping trials (2 − 3), this would amount to a POD of two. We divided PODs by the maximum distance (4), normalizing the score between zero and one:

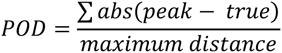

On this distance metric, lower values indicate a closer correspondence between ping-induced peaks and condition presentation order, which in turn confers stronger evidence for ping-based decoding enhancement. For our statistical evaluation, we used a two-level permutation approach (similar to van Bree et al., 2022). Specifically, we compared the distribution of empirical PODs with PODs calculated across 10^6^ second-level permutations, randomly grabbing from the pool of first-level shuffled decoder time courses. The p-values were defined by the resulting percentile of the empirical POD within the distribution of second-level shuffled PODs (one-sided test, empirical < permuted).

For the detection of decoder peaks in this analysis, we detected the maximum peak in the derivative of the cumulative sum of decoding time series. We chose this peak detection method over more standard approaches—such as simply extracting the largest peak from raw decoding series— because independent simulations revealed that this algorithm is most powerful at detecting true POD effects, outperforming a range of competing approaches (Supplementary Materials; Section 2).

## Results

### Behavioural results

As expected in light of our experimental design, participants achieved high memory recognition performance, with scores approaching ceiling across behavioural analyses. First, we found no significant difference in memory performance across participants between the ping (M = 0.980, SE = 0.0032) and no ping condition (M = 0.984, SE = 0.006) during the recognition phase (t(27) = -0.745, p = 0.463; Fig. 1B), suggesting that the decoding analyses that follow are not influenced by absolute inter-condition differences in behaviour. This general near-ceiling performance is also apparent when analysing recognition performance across participants (M = 0.980, SD = 0.015; Fig. 1C) and blocks (M = 0.980, SD = 0.009; Fig. 1D). Furthermore, participants reported a high rate of remembered to forgotten judgments during the recall phase (M = 0.819; SD = 0.022). The average RT during encoding was 2313 ms for subsequently recognized trials (SD = 1041 ms; n = 2194 trials), and 2472 ms for subsequently forgotten trials (SD = 1105 ms; n = 46 trials; Fig. 1E).

### Event-related potentials

We observed a robust evoked EEG response after pings (Fig. 2). Specifically, for each of the three stimulus onset asynchrony (SOA) conditions, we observed an extended peak of activity across occipitoparietal channels that followed the distribution of ping times for retrieval cue-locked data, peaking approximately 200 to 300 ms after pings. To further confirm that pings successfully evoked a visual response, we applied a ping-locked analysis across all channels and found significantly higher ERP amplitudes after pinged than no-pinged trials in posterior channels (Fig. 2, insets). Together, the ERP analysis suggests pings yielded a strong time-locked response that could putatively interact with ongoing LTM representations. For cue-locked and ping-locked ERPs for each participant, time-resolved topographical plots, and for p-values of each channel in Fig. 2 inset topographies, see the Supplementary Materials (Section 1).

**Figure 2.**
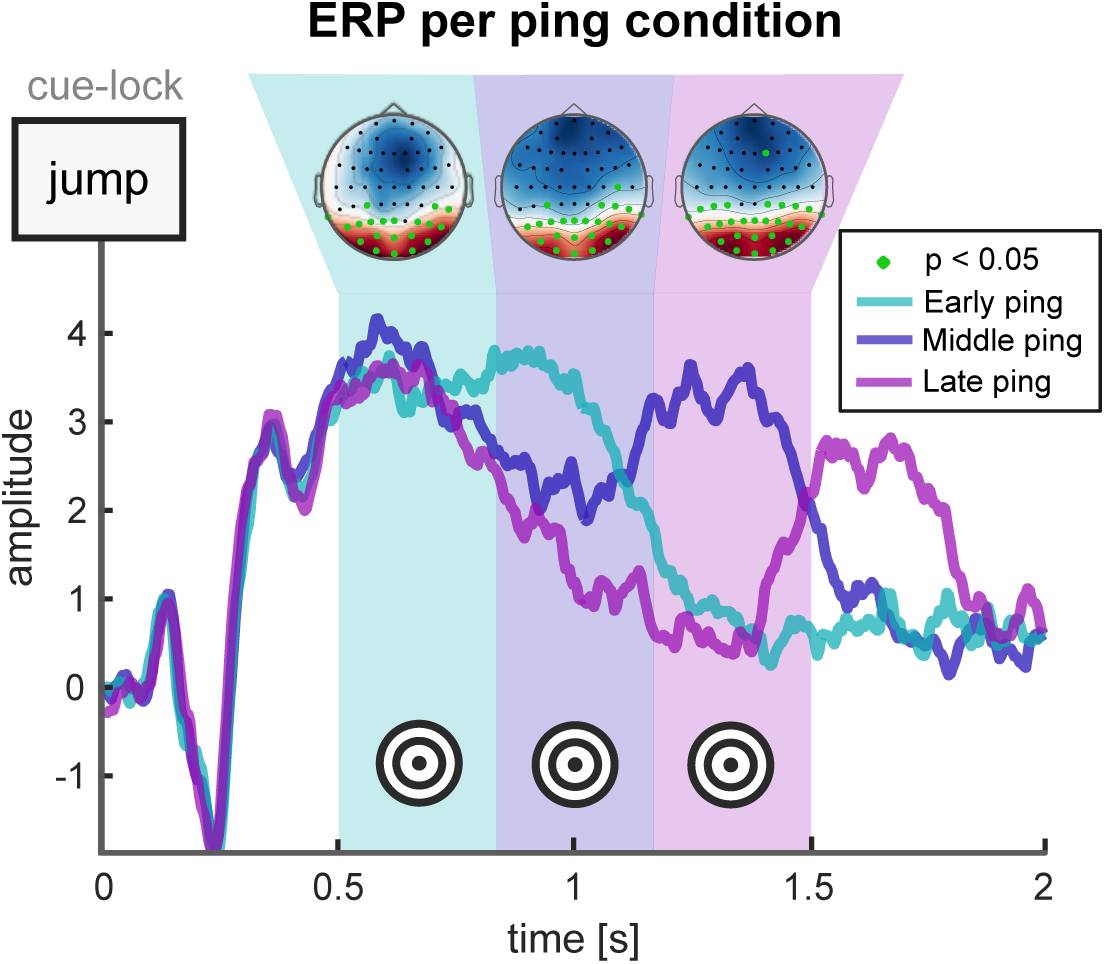
Ping-induced event-related potential. Average evoked response in posterior EEG channels across early (turquoise), middle (blue), and late ping (purple) trials during the recall phase. The inset topographies reveal higher posterior amplitudes following ping trials as contrasted with no-ping trials (Monte Carlo permutation test; Bonferroni-corrected).

### Decoding results

#### Stimulus category selection

We used a multi-class LDA on no-ping trials (25% of the overall recall trials) to determine which retrieved stimulus category (top, middle, or bottom level) is most robustly decodable when our main experimental pinging manipulation was not present (Fig. 3). We found that none of the three levels displayed significant windows of decodability during our retrieval period of interest from 500 to 2000 ms after cue onset (Wilcoxon signed-rank test; p > 0.11 for top; p > 0.25 for middle; p > 0.07 for bot). We proceeded with the top-level, which with its two classes (objects and scenes) afforded simple binary classification with comparatively low variability in decoding performance. Next, during our main analysis, we investigated whether pings enhance the decodability of LTM contents.

**Figure 3.**
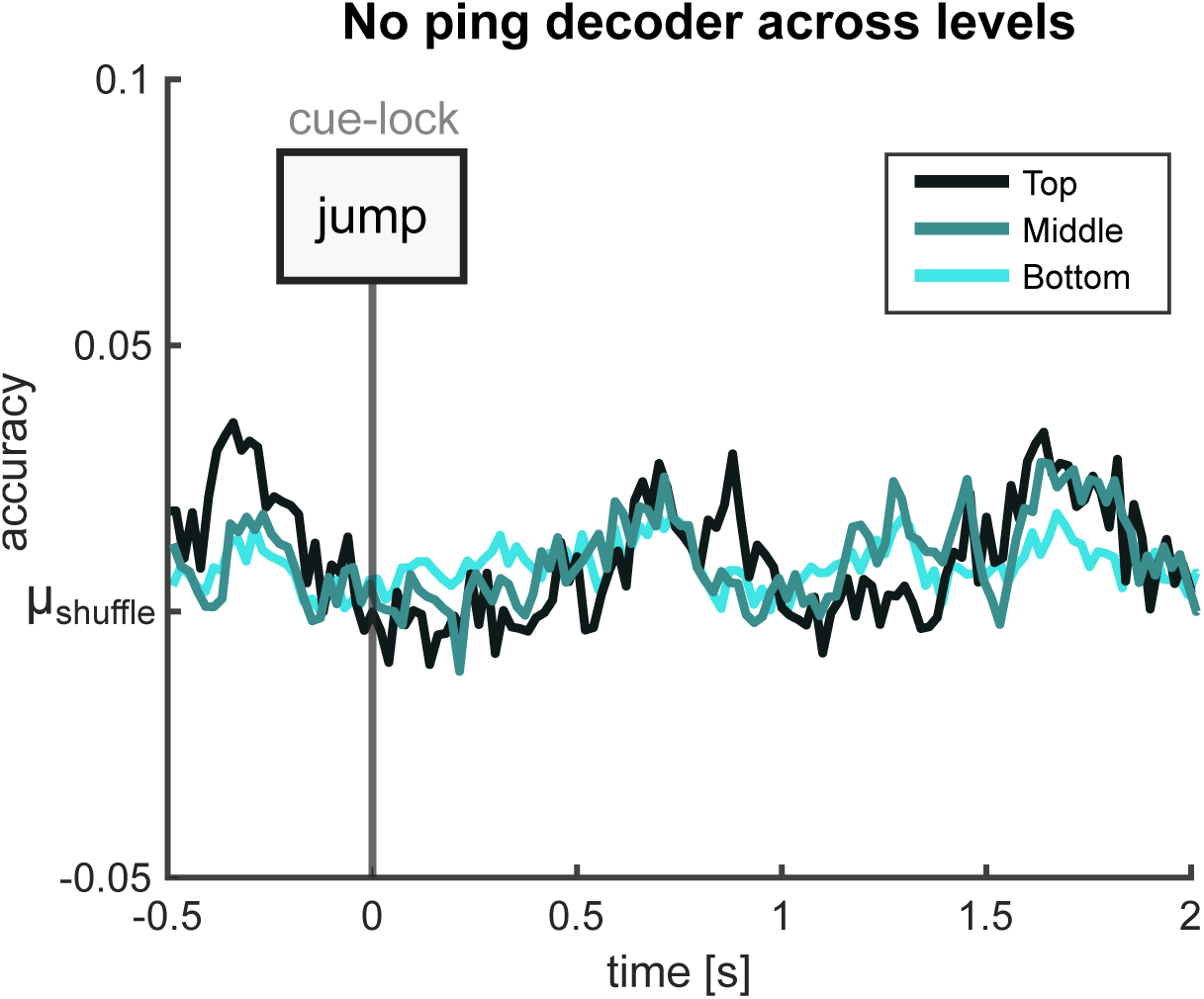
Stimulus category selection. Average decoding accuracy across stimulus category levels (top, middle, bottom). Decoding accuracy was quantified relative to the average performance across shuffled decoding results. No significant differences were observed for any level (Wilcoxon signed rank test, controlled for multiple comparisons using FDR).

#### Main analysis

For our central analysis, we compared decoder performance between ping and no-ping trials for top-level (objects vs scenes) classification, both with the data locked to retrieval cues, and to pings/pseudo-pings (i.e., artificial markers derived from the pool of ping timings; Fig. 4). For the cue-locked analysis, we found no windows where decoding was above chance for no-ping trials (two-level Monte Carlo permutation; p > 0.49; Fig. 4A), while the ping trials showed several significant windows of content decodability (p < 0.05; Fig. 4B). To validate our analysis we carried out a direct comparison between the ping and no-ping trial decoder, as opposed to contrasting each condition with a shuffled baseline. In this analysis, we found no evidence for a ping-induced decodability enhancement; neither in the cue-locked (Wilcoxon signed-rank test; p > 0.99; Fig. 4C) nor in the (pseudo-)ping-locked data (p > 0.99; Fig. 4D).

**Figure 4.**
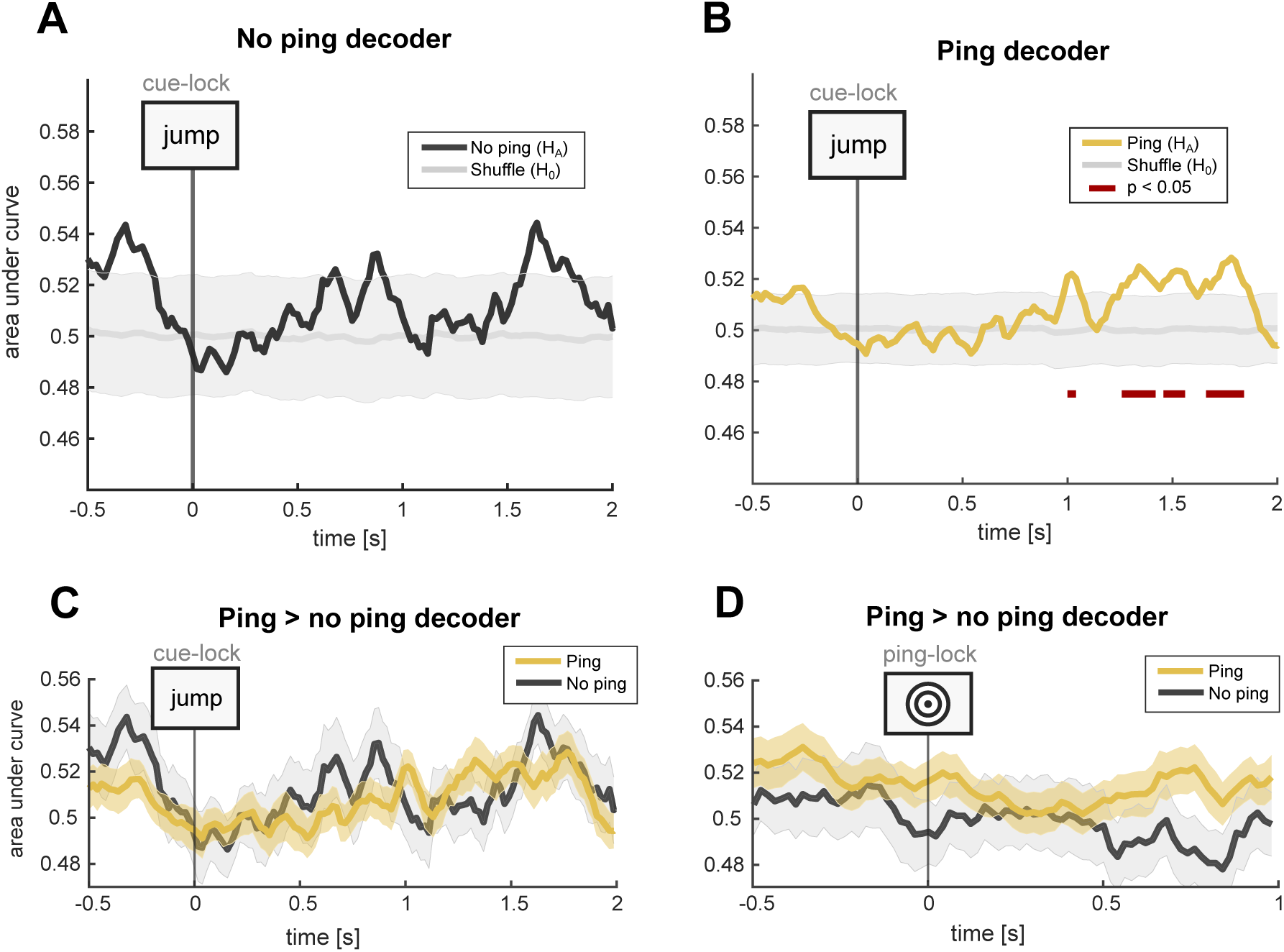
Main decoder analysis. **(A)** Cue-locked decoding across no-ping trials compared with a shuffled baseline. **(B)** Cue-locked decoding across ping trials compared with a shuffled baseline. **(C)** Direct comparison between on ping and no-ping trials. **(D)** Same as (C), but with the data time-locked to pings and (artificially marked) pseudo-pings. In (A) and (B) the shaded area represents the 5^th^ and 95^th^ percentile of the distribution of 2^nd^-level permutations of the shuffled decoder, and in (B) and (C) it represents the SEM of the empirical decoder. In (A) and (B), p-values were derived using two-level Monte Carlo permutations, and in (C) and (D) using Wilcoxon signed-rank test (all p-values were corrected using FDR).

In light of an important methodological observation, we place more importance on the latter analysis, which directly compares the empirical decoding performance for ping and no-ping conditions without leveraging shuffled results. Specifically, we observed that the standard error of the mean (SEM) of the shuffled distributions varies substantially between ping (μ_SEM_ = 0.047) and no-ping (μ_SEM_ = 0.028), which we speculated could be explained by trial number differences alone. We inferred that since the ping trial decoder was trained and tested on three times more trials than the no-ping trial decoder, this might naturally shrink SEM values of the shuffled distribution and thereby modulate test statistics. In support of this interpretation, we built a simulation which confirms that an increase in the number of trials (and the number of decoding classes) reduces p-values, but only if there is an effect in the data (Supplementary Materials; Section 3). Therefore, instead of relying on ping-to-shuffle and no-ping-to-shuffle comparisons where power differences might misleadingly lead us to infer a ping-related enhancement, we placed most credence in the direct comparison between ping and no-ping trials in which shuffled results are sidestepped (Fig. 4C & Fig. 4D; see the Supplementary Materials for an extended discussion; Section 3.3).

#### Condition-relative decoding peaks

Next, we turn to the presumably more sensitive peak-order analyses. Qualitatively, we observe no ordered structure in decoder peaks when averaging across participants for each SOA pinging condition (Fig. 5A). For a quantitative analysis, we formally compared peak order structure by comparing POD scores for the empirical and shuffled decoder using two-level permutation tests. This analysis confirmed the previous result by revealing no significant evidence for the hypothesis that pings induce systematic differences in the order of decoding peaks (p = 0.357; Fig. 5B).

**Figure 5.**
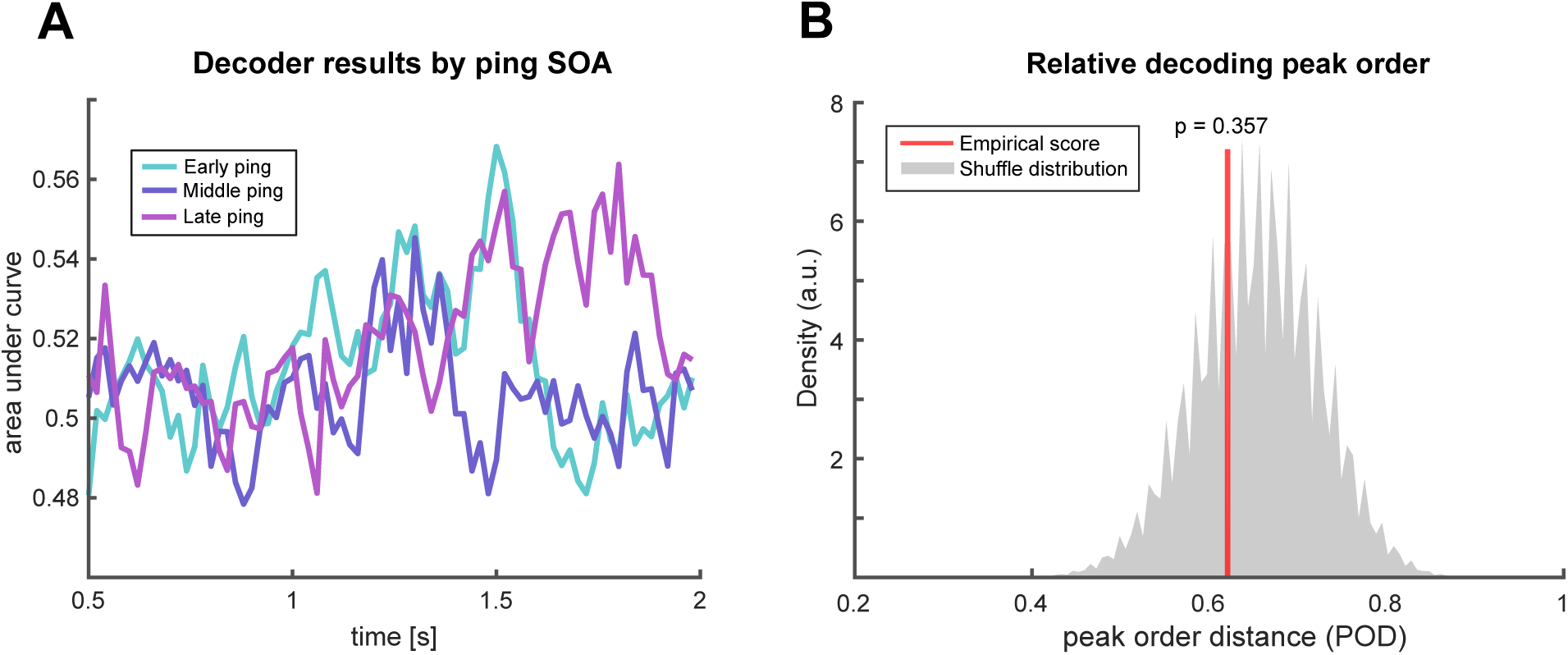
Condition-relative peak analysis. **(A)** Decoding results specific to for early (cyan), middle (blue), and late (purple) ping conditions, averaged across participants. **(B)** Peak order distance scores for the empirical decoder (red line) among a pool of 2^nd^-level permutations derived from the shuffled decoder (grey distribution).

## Discussion

In this study, we set out to systematically evaluate visual perturbation, or ping-based stimulation, as a method to dynamically enhance the decodability of reactivated neural representations during memory recall. Such an approach could supplement offline analytical approaches by adding further read-out enhancements online at the experiment side. Despite promising results in the WM literature, in this LTM context we found no evidence for a ping-based enhancement across several time-resolved decoding analyses. While pings evoked a strong brain response, they did not detectably boost neural signatures of memory representations in EEG data. We draw this conclusion based on two key results. First, in the main comparison between pinged trials and non-pinged trials, we found no significant decoding difference regardless of whether the data was locked to (pseudo-)pings or retrieval cues. Second, in a more advanced analysis that leverages the constraining information of ping presentation timings during the experiment, we also found no evidence for ping-related decoding increases.

There are three overarching explanations for these null results. First, there could be an effect in the data that was left undetected analytically or statistically. Second, there could be an effect that manifests across other experimental contexts, but not with this study’s parameters. Third, there could be no effect in principle, with LTM-based retrieval eluding the enhancement of representational readouts using pings. We consider each option in turn.

First, the signal analysis parameter space is high, with variability in parameters across preprocessing and statistical analysis steps potentially altering the results. One important source of variability concerns the implementation of decoding techniques. Namely, we do not rule out that untested decoding methods such as linear approaches beyond LDA or non-linear classifiers would have resulted in performance enhancements induced by pings. More trivially, our analyses could have been optimal, with our key statistical results containing a type-II statistical error.

Second, the parameter space on the experimental side is also high. Here, we opted for a word-image association task, which has previously been shown to afford classification-based inferences about memory processing in the brain (Linde-Domingo et al., 2019; Martín-Buro et al., 2020; Mirjalili et al., 2021; Kerrén et al., 2022). However, other LTM tasks might be better suited to reveal ping-based enhancements. Besides the memory task itself, a key set of parameters concerns the presentation of pings. In this study, we chose a high-intensity, short-lasting ping presented with a uniform distribution between 500 and 1500 ms after retrieval cues. This time window was selected based on a review of the timeline of memory reactivation during cued recall, which suggested a maximal content reinstatement within this period (Staresina & Wimber, 2019). However, we observed that decoding was highest late within and even after this range, at approximately 1200 – 2000ms after cue (see Fig. 4D). Decoding plateaus that exceed 1500ms have also been observed in recent work that employed a similar task and analysis pipeline (Kerrén et al., 2022). This raises the possibility that the aforementioned 500 to 1500 ms window is biased to be too early—perhaps because it was estimated based on intracranial EEG research where recordings tend to focus on the hippocampus and other regions that activate early during retrieval (Merkow et al., 2015; Mormann et al., 2005; Staresina et al., 2019). Put differently, it is possible that we did not find significant effects because the signatures of retrieved contents tended to arise robustly only after our ping presentation times. We recommend that future work considers later ping times, potentially informed by maximum decodability periods found in this and other work, or ideally in newly acquired pilot data. Moreover, additional research could explore parameters such as ping duration, intensity, and strength. Furthermore, besides visual pings, a plethora of other perturbational approaches are on stock that could realize the ping’s proposed effects. Also inspired by WM research, stimulation using auditory impulses might offer a multimodal route to improving the readout of LTM contents (Kandemir & Akyürek, 2023). Furthermore, brain stimulation methods like transcranial magnetic and ultrasound stimulation have the potential to regularize brain activity through the induction of a dynamics-altering magnetic or ultrasound pulse (Moliadze et al., 2003; Mueller et al., 2014).

A third possibility is that none of these factors explain our null results, with ping-based approaches restricting their utility to WM tasks. One specific possibility could be that WM and LTM differ in their mechanisms of action, with separate kinds of neural processes underpinning them. Indeed, classically WM is believed to involve the active maintenance of stimulus-induced information (Fuster & Alexander, 1971; Goldman-Rakic, 1995), whereas LTM is assumed to be based on a generative reconstruction of past experience based on the activation of silent information-storing engrams (Josselyn & Tonegawa, 2020). Perhaps the sweep of activity associated with the ping interacts more effectively with functional brain activity maintained continuously from stimulus onset, thus explaining WM-to-LTM differences. Speaking against this interpretation is work that suggests WM representations are encoded in activity-silent networks through short-lasting synaptic changes (Kamiński & Rutishauser, 2020; Masse et al., 2020; Stokes, 2015), which would not be fundamentally different from how LTM works. Contradicting this in turn is a critique which argues that evidence for activity-silent networks in WM tasks could alternatively be explained by LTM processes kicking in (Beukers et al., 2021). Thus, since it is both unclear to what extent the mechanisms of WM and LTM differ and to what extent WM and LTM intertwine in studies where ping-based effects have been demonstrated, we avoid firm interpretations in this part of the possibility space. In summary, although pings unambiguously elicited expected patterns of visual activity (Fig. 2), we failed to find effects on memory decoding, either because they were left undetected in our analysis, because they do not show up in our experimental protocol, or because they do not exist.

This study builds on decoding research that investigates the physical basis of memory, leveraging its findings for a strictly instrumental purpose: the systematic enhancement of LTM readouts. This undertaking is key because the field presently lacks temporally sensitive neuroimaging methods that enable the consistent and clear readout of memory representations, which is needed to explain how the brain implements memory processes. Furthermore, the analytical challenges, null results, and possible solutions considered in this work could inform practice in fields closely aligned with memory, such as the neuroscience of mental imagery (Dijkstra et al., 2018).

To conclude, most efforts to improve memory readouts from electrophysiology data have been restricted to the signal analysis end. Here, we advocate for research that explores online manipulations as memory tasks are unfolding, which has previously shown to complement or synergize with decoding techniques. For long-term memory decoding in particular however, such interventions are scarce, which limits research because memory involves low decodability to begin with. Thus, even if a further carving out of the parameter space does not demonstrate a notable benefit of visual perturbations, future research should creatively explore alternative online methods such as multimodal stimulation and non-invasive brain stimulation.

## Acknowledgments

We thank David Rose, Janvi Sidhu, and Jacqueline McDiarmid for their assistance during data acquisition. This work was supported by a Starting Grant from the European Research Council awarded to MW (ERC-2016-StG-715714).

## Supplementary Materials

### Event-related potential

**Supplementary Figure 1.**
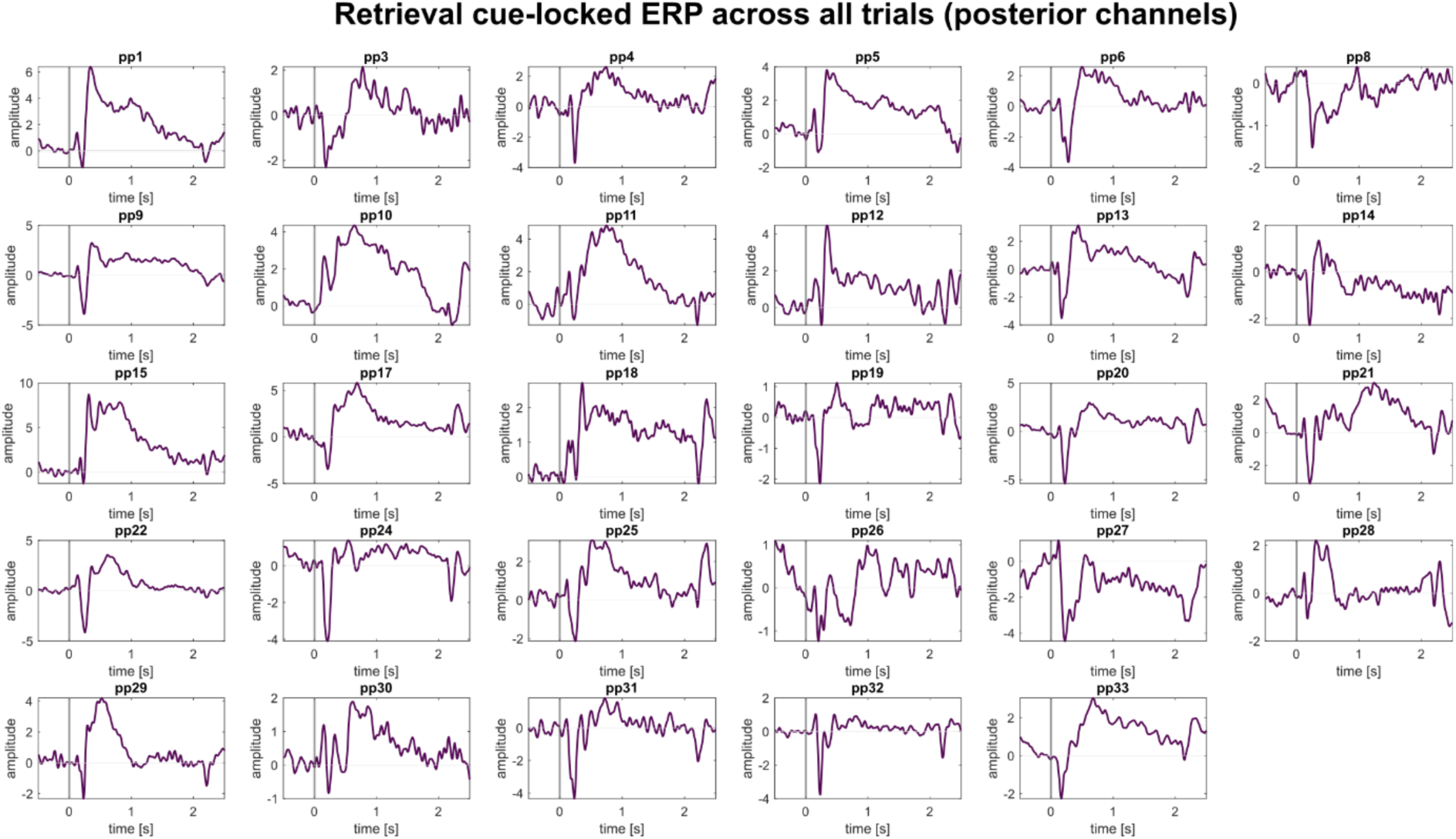
Retrieval cue-locked ERP. The purple trace reflects the average cue-locked response for each participant across posterior EEG channels. The grey horizontal line represents cue onset. For more details, see the Methods section in the main text. The amplitude on the y-axis is in arbitrary units.

**Supplementary Figure 2.**
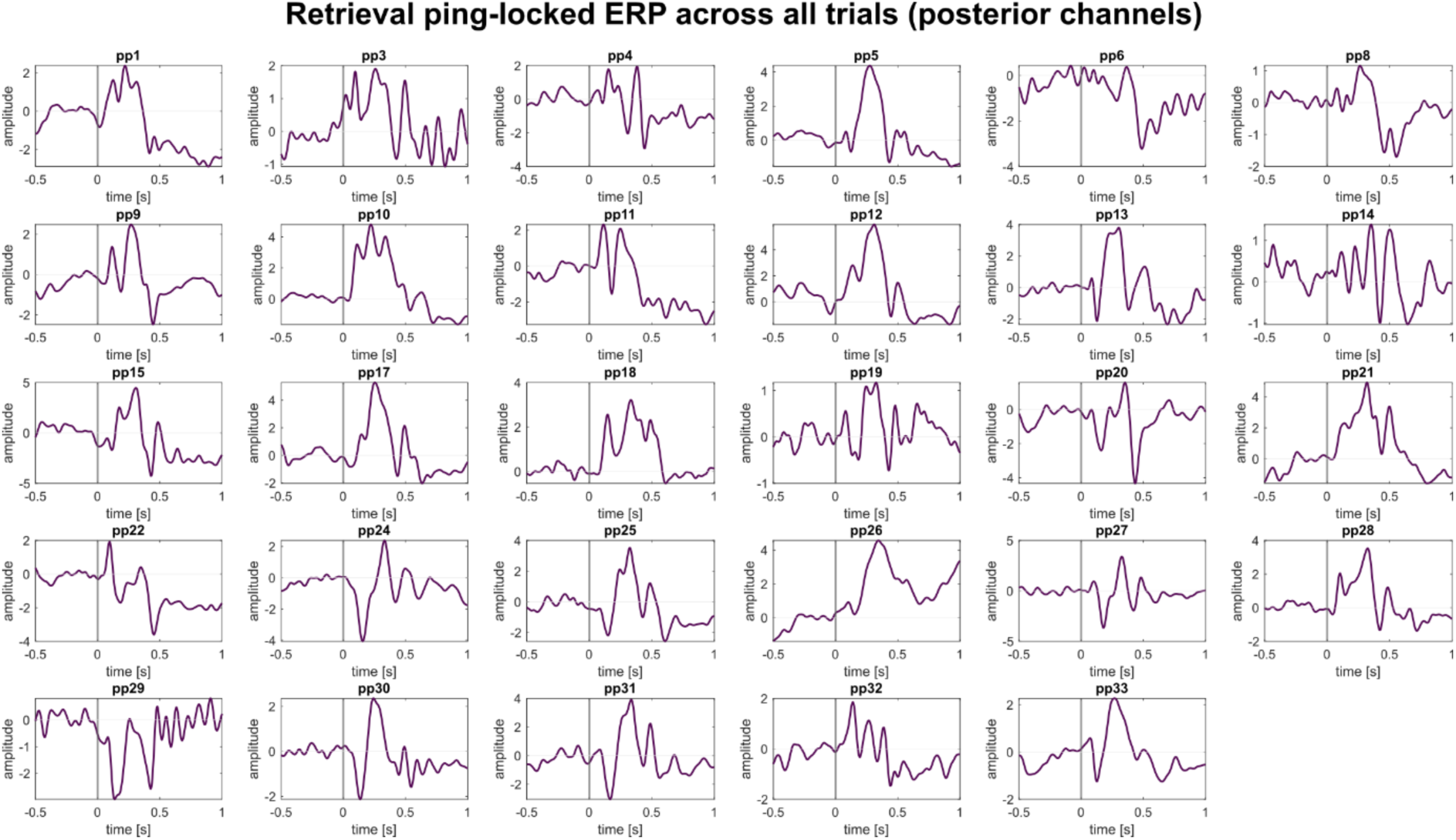
Retrieval ping-locked ERP. The purple trace reflects the average ping-locked response for each participant across posterior EEG channels. The grey horizontal line represents ping onset. For more details, see the Methods section in the main text. The amplitude on the y-axis is in arbitrary units.

**Supplementary Figure 3.**
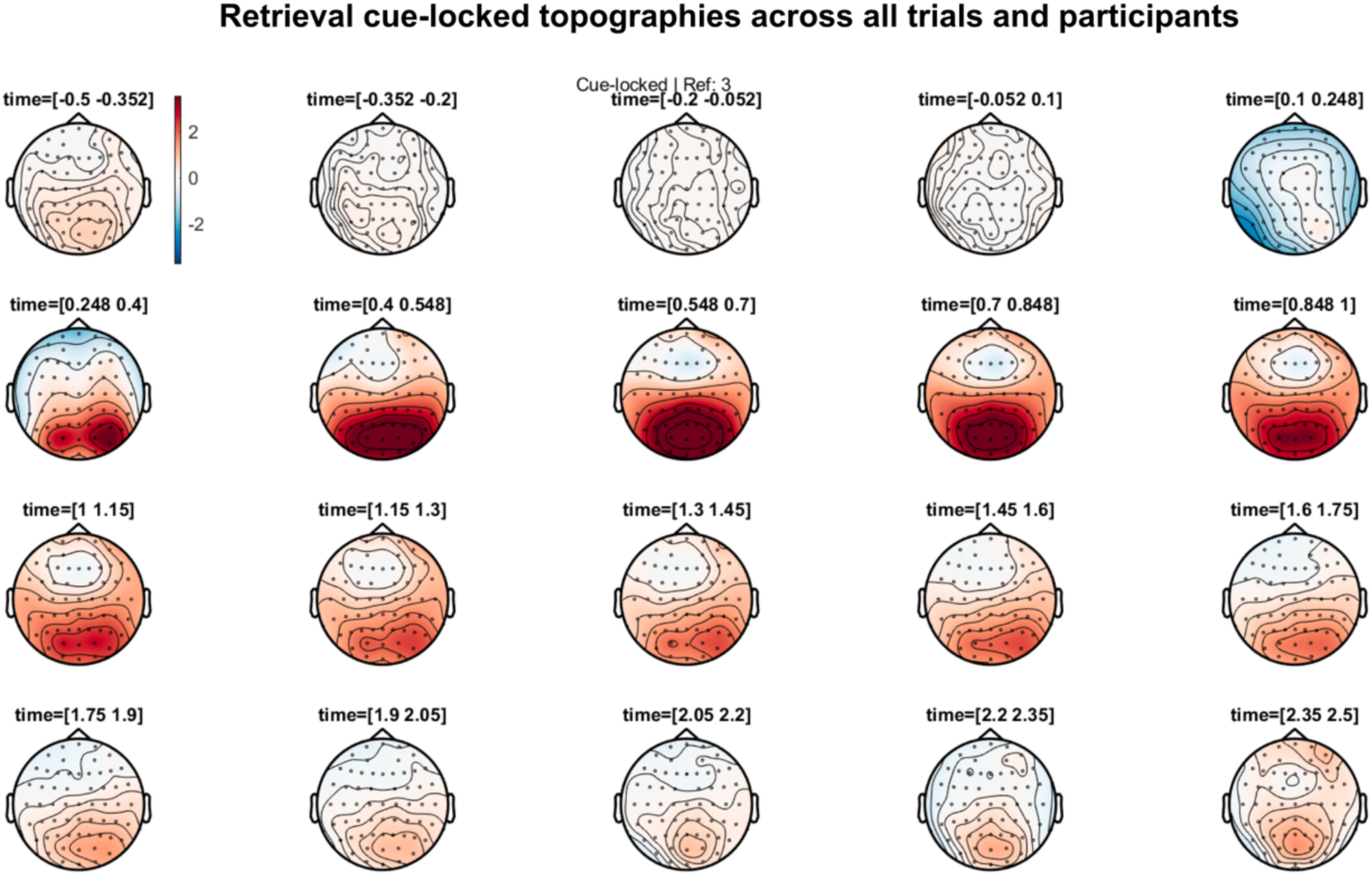
Retrieval cue-locked topographies. These topographical plots represent the average cue-locked activity across participants. The colours represent the difference in EEG activity before and after cue onset in arbitrary units (red colours represent activity_post_ > activity_pre_ and vice versa for blue colours). No statistical analysis was carried out for these topographical contrasts. For more details, see the Methods section in the main text.

**Supplementary Figure 4.**
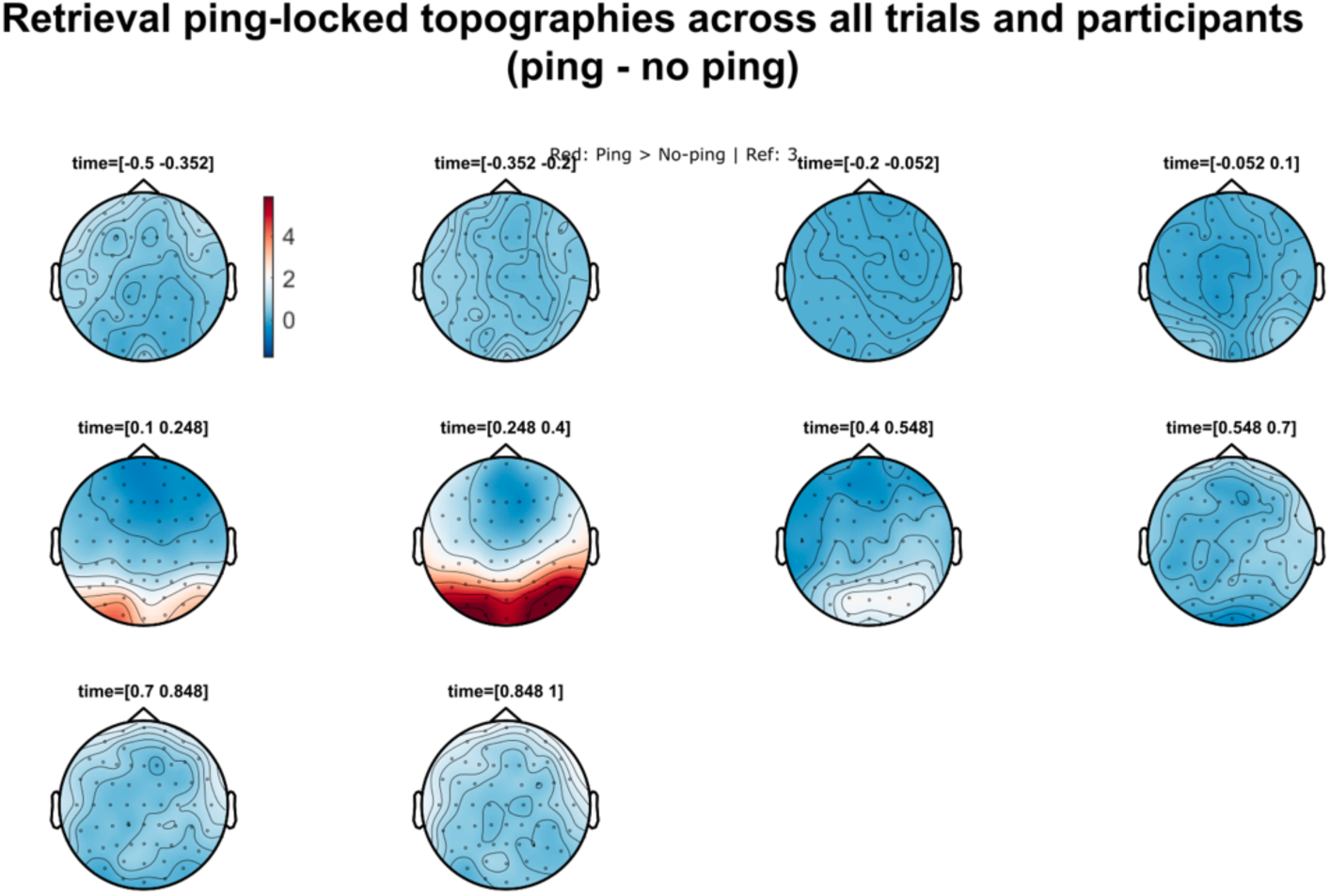
Retrieval ping-locked topographies (ping vs. no ping trials). These topographical plots represent the average cue-locked activity across participants. The colours represent the difference in EEG activity between ping and no-ping (red colours represent activity_ping_ > activity_no ping_ and vice versa for blue colours). No statistical analysis was carried out for these topographical contrasts. For more details, see the Methods section in the main text.

**Supplementary Table 1.**
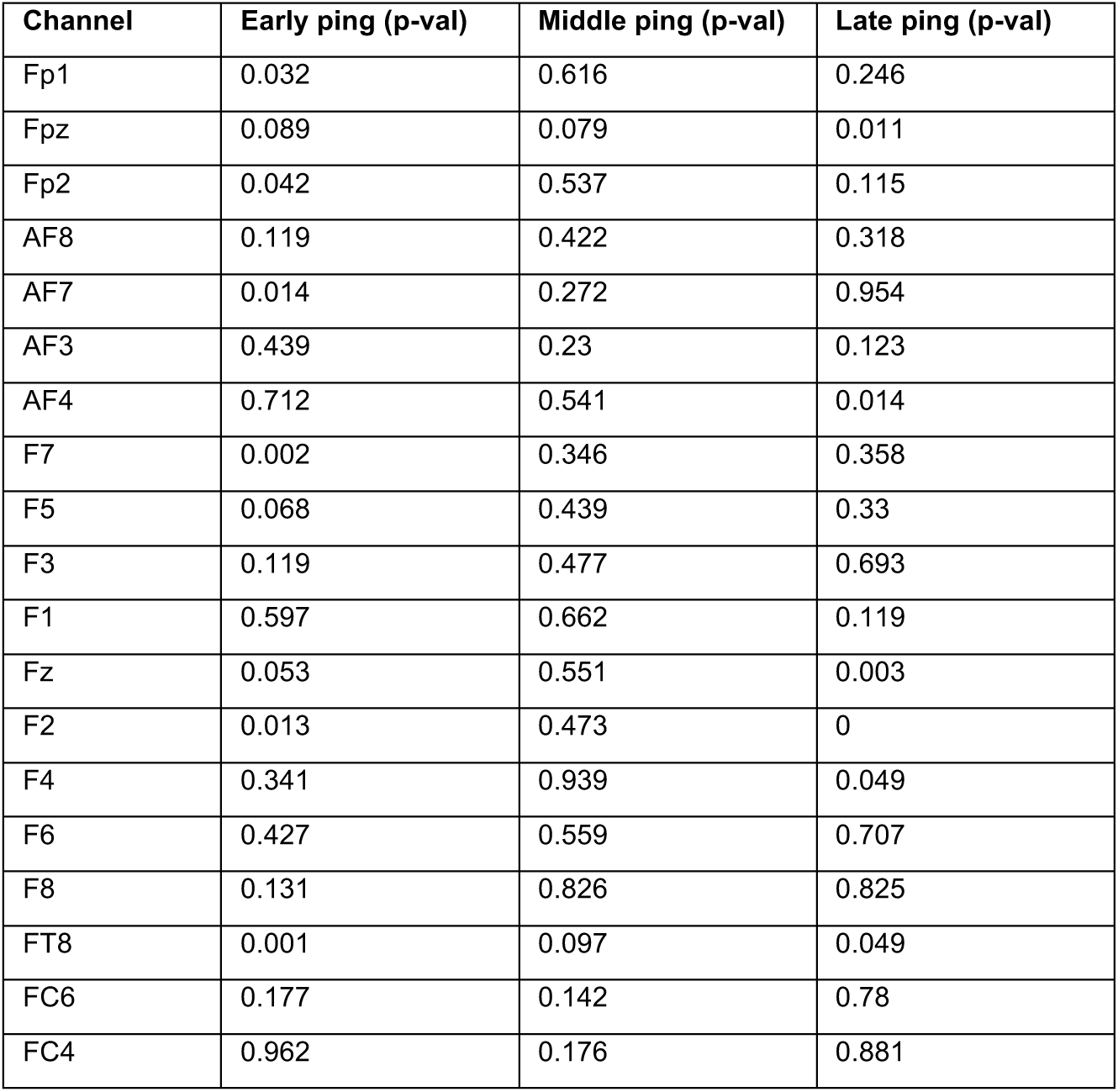
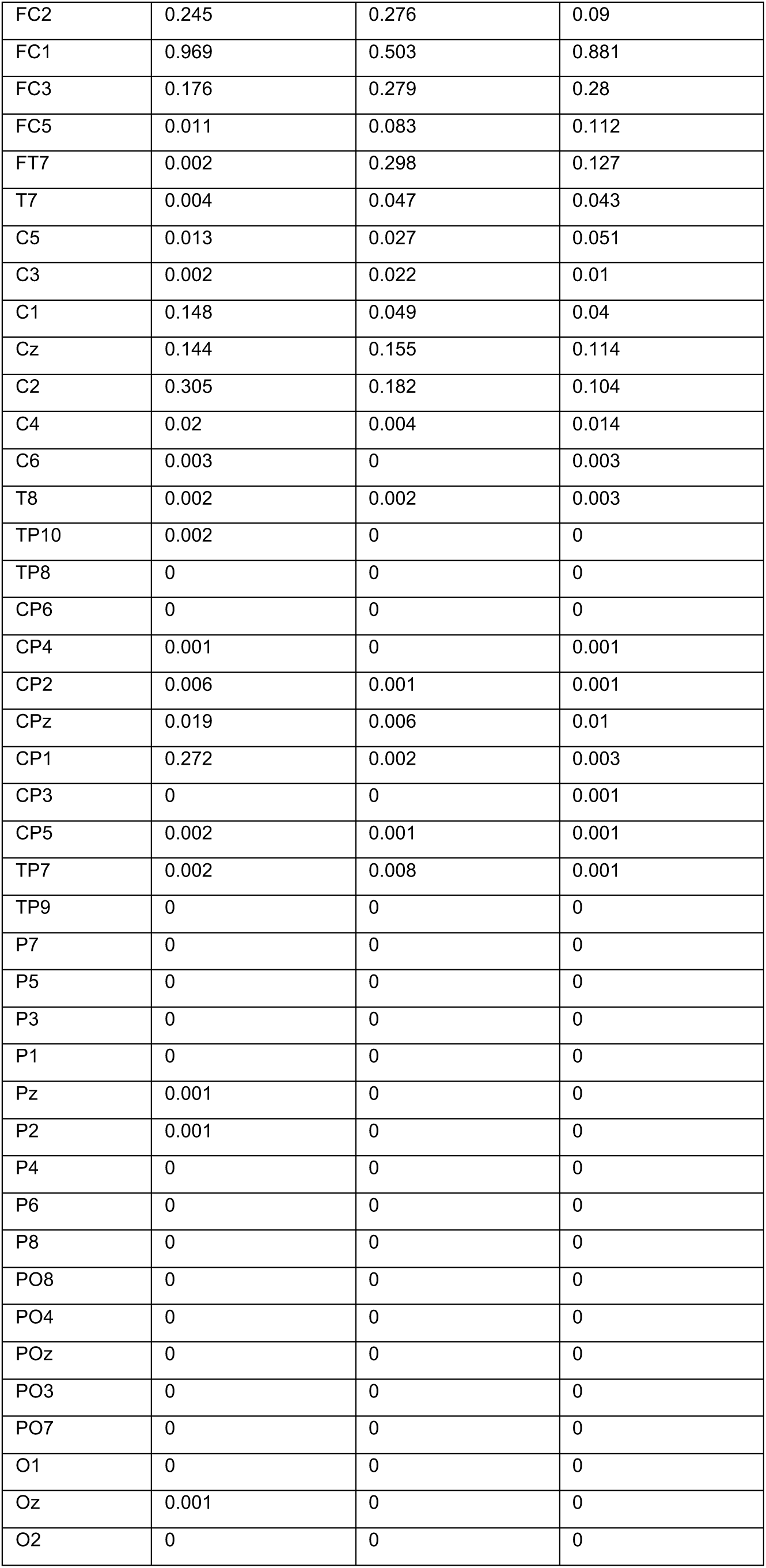
P-values associated with inset topographies in main text Fig. 2; rounded to three decimal points.

| Channel | Early ping (p-val) | Middle ping (p-val) | Late ping (p-val) |
| --- | --- | --- | --- |
| Fp1 | 0.032 | 0.616 | 0.246 |
| Fpz | 0.089 | 0.079 | 0.011 |
| Fp2 | 0.042 | 0.537 | 0.115 |
| AF8 | 0.119 | 0.422 | 0.318 |
| AF7 | 0.014 | 0.272 | 0.954 |
| AF3 | 0.439 | 0.23 | 0.123 |
| AF4 | 0.712 | 0.541 | 0.014 |
| F7 | 0.002 | 0.346 | 0.358 |
| F5 | 0.068 | 0.439 | 0.33 |
| F3 | 0.119 | 0.477 | 0.693 |
| F1 | 0.597 | 0.662 | 0.119 |
| Fz | 0.053 | 0.551 | 0.003 |
| F2 | 0.013 | 0.473 | 0 |
| F4 | 0.341 | 0.939 | 0.049 |
| F6 | 0.427 | 0.559 | 0.707 |
| F8 | 0.131 | 0.826 | 0.825 |
| FT8 | 0.001 | 0.097 | 0.049 |
| FC6 | 0.177 | 0.142 | 0.78 |
| FC4 | 0.962 | 0.176 | 0.881 |
| FC2 | 0.245 | 0.276 | 0.09 |
| FC1 | 0.969 | 0.503 | 0.881 |
| FC3 | 0.176 | 0.279 | 0.28 |
| FC5 | 0.011 | 0.083 | 0.112 |
| FT7 | 0.002 | 0.298 | 0.127 |
| T7 | 0.004 | 0.047 | 0.043 |
| C5 | 0.013 | 0.027 | 0.051 |
| C3 | 0.002 | 0.022 | 0.01 |
| C1 | 0.148 | 0.049 | 0.04 |
| Cz | 0.144 | 0.155 | 0.114 |
| C2 | 0.305 | 0.182 | 0.104 |
| C4 | 0.02 | 0.004 | 0.014 |
| C6 | 0.003 | 0 | 0.003 |
| T8 | 0.002 | 0.002 | 0.003 |
| TP10 | 0.002 | 0 | 0 |
| TP8 | 0 | 0 | 0 |
| CP6 | 0 | 0 | 0 |
| CP4 | 0.001 | 0 | 0.001 |
| CP2 | 0.006 | 0.001 | 0.001 |
| CPz | 0.019 | 0.006 | 0.01 |
| CP1 | 0.272 | 0.002 | 0.003 |
| CP3 | 0 | 0 | 0.001 |
| CP5 | 0.002 | 0.001 | 0.001 |
| TP7 | 0.002 | 0.008 | 0.001 |
| TP9 | 0 | 0 | 0 |
| P7 | 0 | 0 | 0 |
| P5 | 0 | 0 | 0 |
| P3 | 0 | 0 | 0 |
| P1 | 0 | 0 | 0 |
| Pz | 0.001 | 0 | 0 |
| P2 | 0.001 | 0 | 0 |
| P4 | 0 | 0 | 0 |
| P6 | 0 | 0 | 0 |
| P8 | 0 | 0 | 0 |
| PO8 | 0 | 0 | 0 |
| PO4 | 0 | 0 | 0 |
| POz | 0 | 0 | 0 |
| PO3 | 0 | 0 | 0 |
| PO7 | 0 | 0 | 0 |
| O1 | 0 | 0 | 0 |
| Oz | 0.001 | 0 | 0 |
| O2 | 0 | 0 | 0 |

### Peak order analysis simulation

#### 2.1 Time series simulation

We used MATLAB (the MathWorks) to generate time series with two components: (1) a peak at a fixed time point (1000 ms), and (2) autocorrelated noise generated using a random walk procedure. We matched several characteristics of the simulated time series to our empirical decoding data, including the analysis period (500 to 2000 ms), sampling rate (50 Hz), and the number of (virtual) participants (N = 29). The signal-to-noise (SNR) ratio of the simulation was set to 1.15, qualitatively matching peaks observed in the empirical data. We found that varying the SNR does not significantly alter the results. We generated 1000 trials per participant, resulting in 29000 trials in total.

#### 2.2 Analysis

We included a smoothing parameter that implemented one of four smoothing methods: no filter, a Gaussian filter, a Savitzky-Golay filter, and a median filter. We also included a window size for smoothing, set to 10 samples for our main analysis. We compared the performance of eight peak detection methods, evaluating each of them based on the absolute distance between estimated peaks and true peaks—amounting to a simplified version of the *peak order distance* score described under *condition-relative decoding peaks* in the main text. The winning method was locked in for our empirical analysis. We tested eight peak detection methods:

1. Low-pass approach, where the maximum peak was computed after a low-pass filter was applied to the time series.
2. Maximum value approach, which simply computed the maximum value per time series regardless of whether the surrounding data was peak-like.
3. Cumulative sum approach, which computed the maximum peak in the derivative of the cumulative sum of the data.
4. Cumulative integral approach, which computed the maximum peak in the cumulative integral of the data via the trapezoidal method.
5. Integral cumulative sum approach, which worked as the previous method but which operates over the cumulative sum rather than raw time series.
6. Wavelet transform-based method, which finds the maximum peak in a wavelet decomposed version of the data.
7. Hilbert transform-based method, which find the maximum peak in the amplitude fluctuations in the envelope of the time series.
8. Cross-correlation method, which finds the time lag with a maximal correlation between the signal and iteratively shifted versions of itself.

#### 2.3 Results

We found that approach 5—the integral cumulative sum approach—reliably achieves low absolute distance errors across parameters (Supplementary Figure 5). These results were generally unchanged across adjustments of the parameters (to evaluate this, we refer to the code published with this manuscript). Thus, we used approach 5 in our main peak order detection analysis.

**Supplementary Figure 6.**
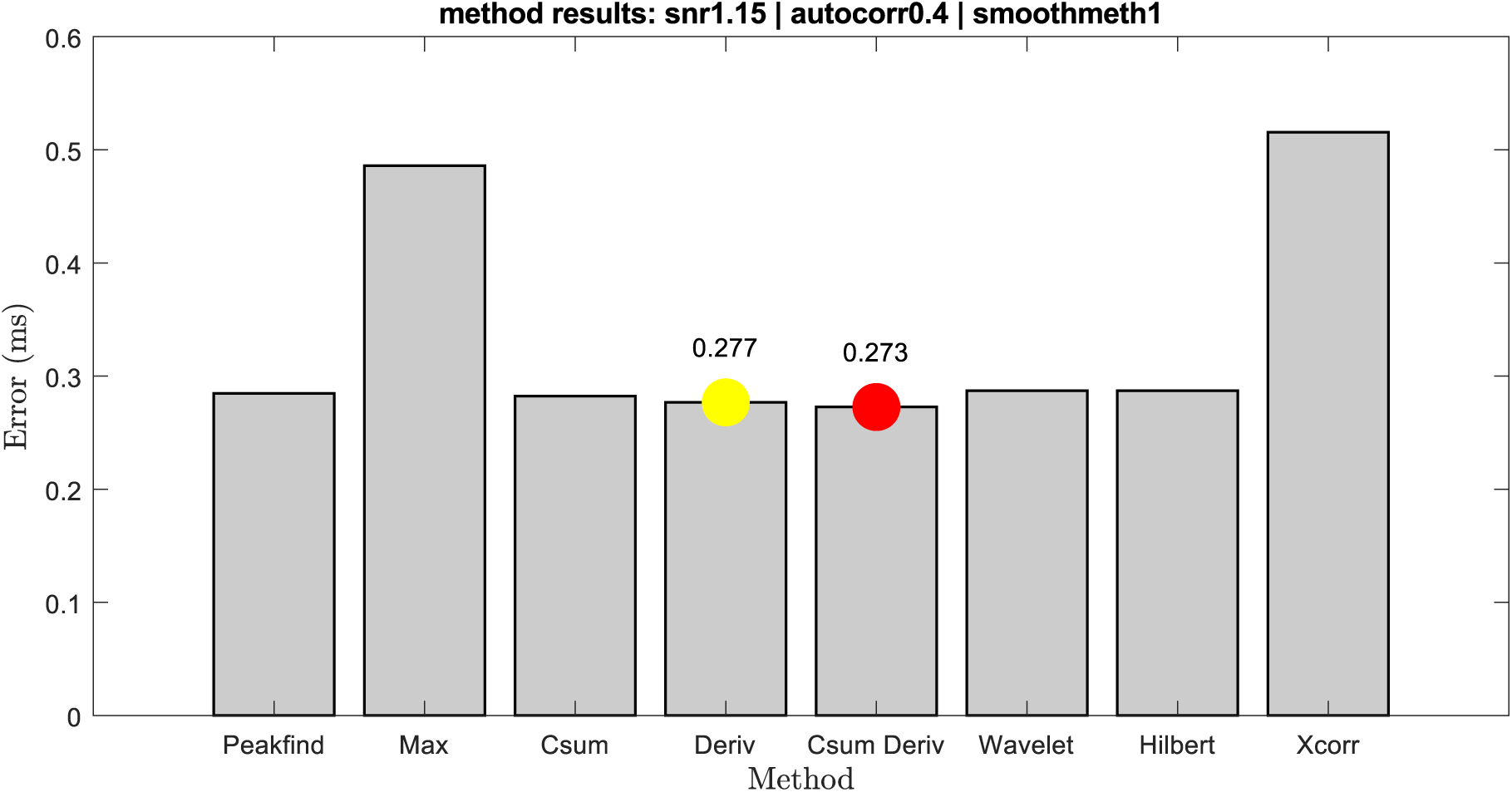
In simulated time series, the integral cumulative sum approach works best for detecting a peak in noisy time series. The red circle indicates the best-performing method, and yellow the second best-performing method. Errors were computed based on the absolute distance in milliseconds (ms) between estimated and true peak location.

### Class and trial number decoding simulation

We speculated based on a qualitative inspection of the empirical decoding results that the number of trials (N_trials_) and classes (N_classes_) reduces the statistical significance of decoding results. We evaluated this intuition by demonstrating using simulations that these two parameters do indeed influence the variance of shuffled and empirical results, which in turn affects p-values but only if there is a true effect in the data.

#### 3.1 Time series simulation

Using MATLAB, we generated one ground truth vector of class labels which represented the true class structure in the simulated data. This vector contained a random sequence of integers randomly grabbed between the interval 1 and N_classes_. For example, with 16 classes, the ground truth pattern might have contained a sequence of [2,7,15,4,13,17] and with 2 classes a sequence of [2,2,1,2,1,2].

Then, to simulate shuffled decoding results, we generated a distribution of random sequences of integers identical to the ground truth procedure, but with newly generated random integers. These random sequences represented shuffled decoding results and were scored based on their average element-wise correspondence to the ground truth pattern—which is how decoding accuracy is normally computed. For example, if the permuted vector is [2,1,2,2,1,1] and the true sequence is [2,2,1,2,1,2], the accuracy would be 50% because half of the class labels correspond to the true structure. Trivially, with increasing repetitions the shuffled distribution will approach chance level predictions of the ground truth pattern (i.e., the expected value is exactly at 1/N_classes_).

Finally, to simulate empirical decoding results, we again generated a distribution of random integers identical to the procedure for shuffled and ground truth decoding results. However, for these data we manually injected between 0% and 60% of the ground truth pattern into the otherwise random vector, effectively modulating decoding accuracy. With 0% of the ground truth injected, there is no statistically detectable difference in accuracy between empirical and shuffled decoding results, because the vectors are equally random. With 60%, the encoding results are substantially more accurate than shuffled results, yielding above chance decoding accuracy.

We simplified our simulation by operationalizing the variable N_trials_ as the number of elements in the vector, allowing us to efficiently investigate how the number of observations influences statistical tests. We also compared N_classes_ = 2 and N_classes_ = 16, which respectively match the number of classes for top- and bottom-level category decoding in our main experiment. Both N_trials_ and N_classes_ were independently manipulated in a 2 ∗ 2 factorial design, allowing us to evaluate the contribution of each variable toward statistical outcomes (as a function of effect size).

#### 3.2 Results

First, with respect to N_classes_, we found that increasing the number of classes reduces the spread of both shuffled and empirical decoding results (Supplementary Figure 7; columns). This happens both if there is no true effect in the empirical data, and when a significant proportion of the ground truth is inserted into the empirical data. Second, we found that N_trials_ similarly reduces the variance of both shuffled and decoding results, both across low and high N_classes_ (Supplementary Figure 7; top and bottom half). Thus, we conclude that both factors modulate the likelihood of finding a significant difference between empirical and shuffled results, but only if there is a true effect in the data. Indeed, as we can glean from the results based on non-existent effects, the distributions of empirical and shuffled will overlap regardless of N_trials_ or N_classes_ (Supplementary Figure 7; left half). In contrast, if there is an effect (60% injected ground truth), both N_trials_ and N_classes_ independently increase the distributional distance between empirical and shuffled accuracy values.

#### 3.3 Discussion

We found that N_trials_ and N_classes_ independently reduce the variance of accuracy results, which will affect statistical tests between empirical and shuffled distributions but only if there is an effect in the data. As suggested in the main text, these findings suggest that statistical analyses that depend on variance comparisons between empirical and shuffled distributions should be interpreted with care if it is done across conditions with varying N_trials_ and N_classes_. With regard to our main analysis for example, the fact that the decoder based on pinged trials yields more significant decodability compared to the decoder based on no-pinged trials should be interpreted with caution because there are differences in N_trials_ between the two conditions that could partially or fully explain this effect. More generally, we found that the condition with more trials or more classes is by default more likely to yield significant p-values—but only if a true effect exist.

**Supplementary Figure 7.**
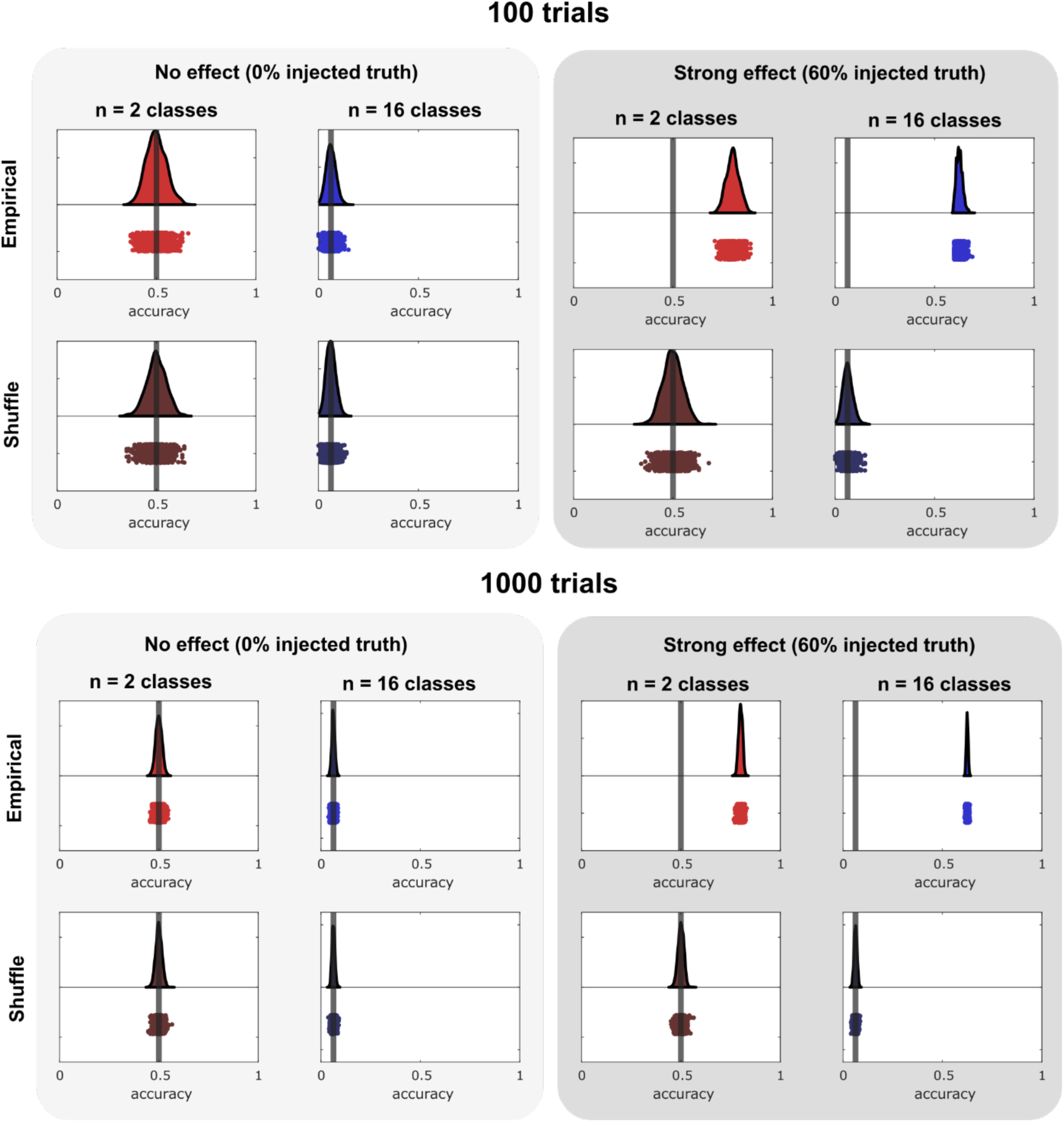
The effects of class and trial number on decoding accuracy. Both the number of classes (columns) and trials (top vs. bottom half) influences the distance between shuffled and empirical distributions—but only if there is an effect in the data (left vs. right half).

These findings may be a manifestation of the classical notion of statistical power in statistical analysis but within the less intuitive context of decoding accuracy. Our interpretation then is not that N_trials_ and N_classes_ must necessarily be equal between conditions for a statistical comparison to be meaningful. Rather, we wanted to err on the side of caution and ensure that analyses where power differences could possibly explain condition differences (e.g., Fig. 3 and Fig. 4A and 4B in the main text) do not inform subsequent analyses and scientific interpretations by themselves. Instead, we supplemented each of the implicated analyses with additional rationale (in the case of Fig. 3) or analyses that do not involve empirical-to-shuffle decoding comparisons. Indeed, Fig. 4C and Fig. 4D involve direct comparisons between empirical and shuffled distributions, sidestepping the issue altogether.

